# Flux-dependent graphs for metabolic networks

**DOI:** 10.1101/290767

**Authors:** Mariano Beguerisse-Díaz, Gabriel Bosque, Diego Oyarzún, Jesús Picóo, Mauricio Barahona

## Abstract

Cells adapt their metabolic fluxes in response to changes in the environment. We present a framework for the systematic construction of flux-based graphs derived from organism-wide metabolic networks. Our graphs encode the directionality of metabolic fluxes via edges that represent the flow of metabolites from source to target reactions. The methodology can be applied in the absence of a specific biological context by modelling fluxes probabilistically, or can be tailored to different environmental conditions by incorporating flux distributions computed through constraint-based approaches such as Flux Balance Analysis. We illustrate our approach on the central carbon metabolism of *Escherichia coli* and on a metabolic model of human hepatocytes. The flux-dependent graphs under various environmental conditions and genetic perturbations exhibit systemic changes in their topological and community structure, which capture the re-routing of metabolic fluxes and the varying importance of specific reactions and pathways. By integrating constraint-based models and tools from network science, our framework allows the study of context-specific metabolic responses at a system level beyond standard pathway descriptions.

## I. INTRODUCTION

Metabolic reactions enable cellular function by converting nutrients into energy, and by assembling macromolecules that sustain the cellular machinery [1]. Cellular metabolism is usually thought of as a collection of pathways comprising enzymatic reactions associated with broad functional categories. Yet metabolic reactions are highly interconnected: enzymes convert multiple reactants into products with other metabolites acting as co-factors; enzymes can catalyse several reactions, and some reactions are catalysed by multiple enzymes, and so on. This enmeshed web of reactions is thus naturally amenable to network analysis, an approach that has been successfully applied to different aspects of cellular and molecular biology, e.g., protein-protein interactions [2], transcriptional regulation [3], or protein structure [4, 5].

Tools from graph theory [6] have previously been applied to the analysis of structural properties of metabolic networks, including their degree distribution [7–10], the presence of metabolic roles [11], and their community structure [12–15]. A central challenge, however, is that there are multiple ways to construct a network from a metabolic model [16]. For example, one can create a graph with metabolites as nodes and edges representing the reactions that transform one metabolite into another [7, 8, 17, 18]; a graph with reactions as nodes and edges corresponding to the metabolites shared among them [19–21]; or even a bipartite graph with both reactions and metabolites as nodes [22]. Importantly, the conclusions of graph-theoretical analyses are highly dependent on the chosen graph construction [23]. A key feature of metabolic reactions is the directionality of flux: metabolic networks contain both irreversible and reversible reactions, and reversible reactions can change their direction depending on the cellular and environmental contexts [1]. Many of the existing graph constructions, however, lead to undirected graphs that disregard such directional information, which is central to metabolic function [8, 16]. Furthermore, current graph constructions are usually derived from the whole set of metabolic reactions in an organism, and thus correspond to a generic metabolic ‘blueprint’ of the cell. However, cells switch specific pathways ‘on’ and ‘off’ to sustain their energetic budget in different environments [24]. Hence, such blueprint graphs might not capture the specific metabolic connectivity in a given environment, thus limiting their ability to provide biological insights in different growth conditions.

In this paper, we present a flux-based approach to construct metabolic graphs that encapsulate the directional flow of metabolites produced or consumed through enzymatic reactions. The proposed graphs can be tailored to incorporate flux distributions under different environmental conditions. To introduce our approach, we proceed in two steps. We first define the *Probabilistic Flux Graph* (PFG), a weighted, directed graph with reactions as nodes, edges that represent supplier-consumer relationships between reactions, and weights given by the probability that a metabolite chosen at random is produced/consumed by the source/target reaction. This graph can be used to carry out graph-theoretical analyses of organism-wide metabolic organisation independent of cellular context or environmental conditions. We then show that this formalism can be adapted seamlessly to construct the *Metabolic Flux Graph* (MFG), a directed, environment-dependent, graph with weights computed from Flux Balance Analysis (FBA) [25], the most widespread method to study genome-scale metabolic networks.

Our formulation addresses several drawbacks of current constructions of metabolic graphs. Firstly, in our flux graphs, an edge indicates that metabolites are produced by the source reaction and consumed by the target reaction, thus accounting for metabolic directionality and the natural flow of chemical mass from reactants to products. Secondly, the Probabilistic Flux Graph discounts naturally the over-representation of pool metabolites (e.g., adenosine triphosphate (ATP), nicotinamide adenine dinucleotide (NADH), protons, water, and other co-factors) that appear in many reactions and tend to obfuscate the graph connectivity. Our construction avoids the removal of pool metabolites from the network, which can change the graph structure drastically [26–30]. Finally, the Metabolic Flux Graph incorporates additional biological information reflecting the effect of the environmental context into the graph construction. In particular, since the weights in the MFG correspond directly to fluxes (in units of mass per time), different biological scenarios can be analysed using balanced fluxes (e.g., from different FBA solutions) under different carbon sources and other environmental perturbations [16, 25, 31, 32].

After introducing the mathematical framework, we showcase our approach with two examples. Firstly, in the absence of environmental context, our analysis of the PFG of the core model of *Escherichia coli* metabolism [33] reveals the importance of including directionality and appropriate edge weights in the graph to understand the modular organisation of metabolic sub-systems. We then use FBA solutions computed for several relevant growth conditions for *E. coli*, and show that the structure of the MFG changes dramatically in each case (e.g., connectivity, ranking of reactions, community structure), thus capturing the environment-dependent nature of metabolism. Secondly, we study a model of human hepatocyte metabolism evaluated under different conditions for the wild-type and in a mutation found in primary hyperoxaluria type 1, a rare metabolic disorder [34], and show how the changes in network structure of the MFGs reveal new information that is complementary to the analysis of fluxes predicted by FBA.

## II. RESULTS

### A. Definitions and background

Consider a metabolic network composed of *n* metabolites *X_i_* (*i* = 1*,…, n*) that participate in *m* reactions

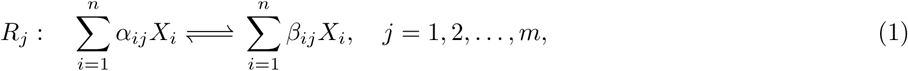

where α_*ij*_ and β_*ij*_ are the stoichiometric coefficients of species *i* in reaction *j*. Let us denote the concentration of metabolite *X_i_* at time *t* as *x_i_*(*t*). We then define the *n*-dimensional vector of metabolite concentrations: **x**(*t*) = (*x*_1_(*t*)*,…, x_n_*(*t*))^*T*^. Each reaction takes place with rate *v_j_*(**x***, t*), measured in units of concentration per time [35]. We compile these reaction rates in the *m*-dimensional vector: **v**(*t*) = (*v*_1_(*t*),…, *v*_*m*_(*t*))^*T*^.

The mass balance of the system can then be represented compactly by the system of ordinary differential equations

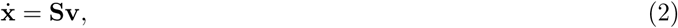

where the *n × m* matrix **S** is the stoichiometric matrix with entries *S_ij_ = β_ij_* − α_*ij*_, i.e., the net number of *X_i_* molecules produced (positive *S_ij_*) or consumed (negative *S_ij_*) by the *j*-th reaction. Figure 1A shows a toy example of a metabolic network including nutrient uptake, biosynthesis of metabolic intermediates, secretion of waste products, and biomass production [32].

**FIG. 1.**
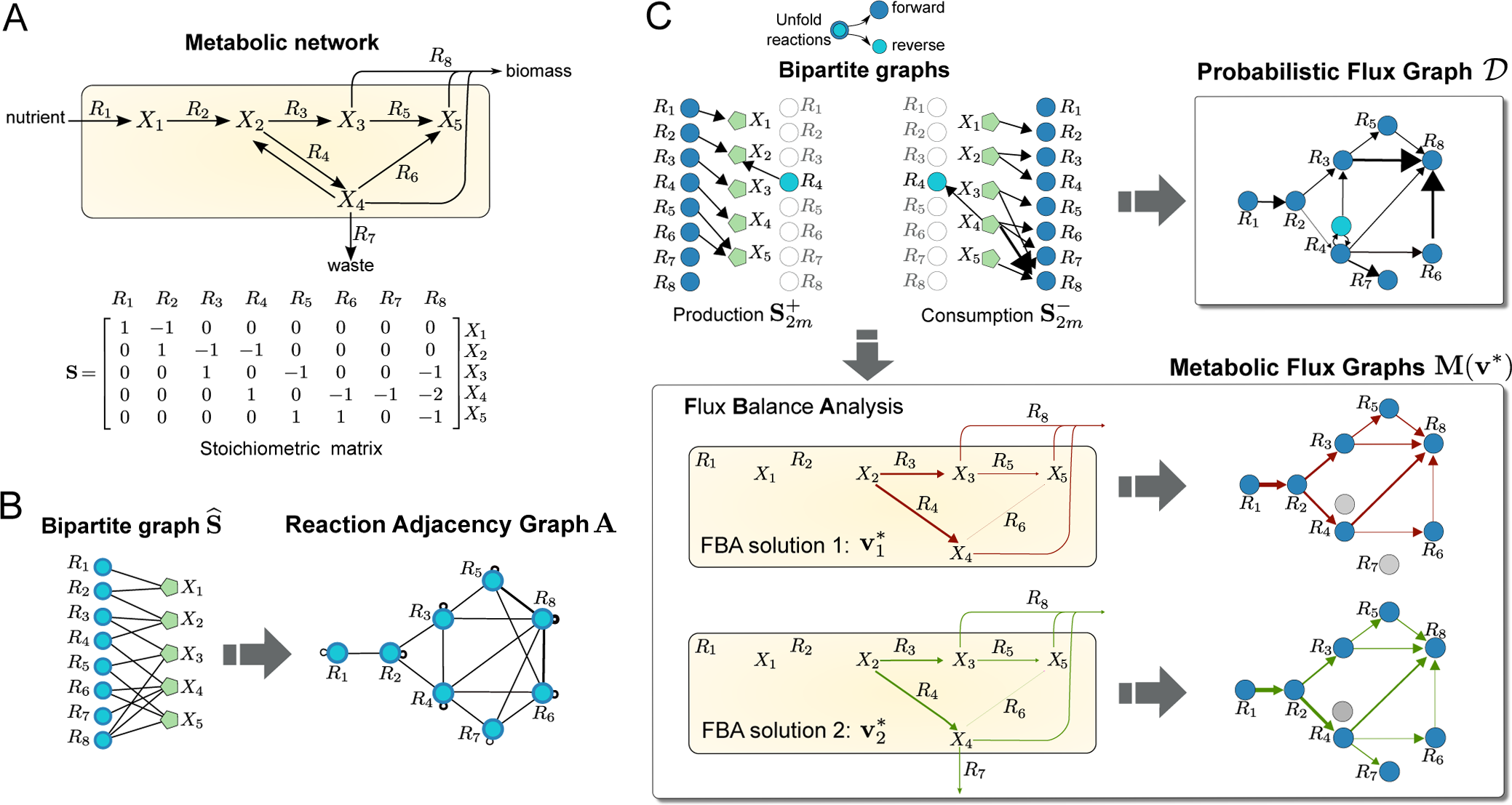
Graphs from metabolic networks. (A) Toy metabolic network describing nutrient uptake, biosynthesis of metabolic intermediates, secretion of waste products, and biomass production [32]. The biomass reaction is *R*_8_ : *X*_3_ + 2*X*_4_ + *X*_5_. (B) Bipartite graph associated with the boolean stoichiometric matrix 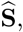, and the Reaction Adjacency Graph (RAG) [16] with adjacency matrix 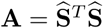. The undirected edges of **A** indicate the number of shared metabolites among reactions. (C) The Probabilistic Flux Graph (PFG) **𝒟** and two Metabolic Flux Graphs (MFG) **M**(**v∗**) constructed from the consumption and production stoichiometric matrices (5). Note that the reversible reaction *R*_4_ is unfolded into two nodes. The PFG in Eq. (8) is a directed graph with weights representing the probability that the source reaction produces a metabolite consumed by the target reaction. The MFGs in (Eq. 12) are constructed from two different Flux Balance Analysis solutions (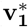 and 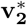) obtained by optimising a biomass objective function under different flux constraints representing different environmental or cellular contexts (see Sec. SI 2 in the Supplementary Information for details). The weighted edges of the MFGs represent mass flow from source to target reactions in units of metabolic flux. The computed FBA solutions translate into different connectivity in the resulting MFGs.

There are several ways to construct a graph for a given metabolic network with stoichiometric matrix **S**. A common approach [16] is to define the *unipartite graph* with reactions as nodes and *m × m* adjacency matrix

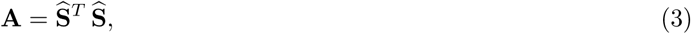

where 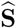 is the boolean version of **S** (i.e., 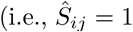 when *S*_*ij*_ ≠ 0 and 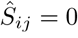 otherwise). This is the *Reaction Adjacency Graph* (RAG), in which two reactions (nodes) are connected if they share metabolites, either as reactants or products. Self-loops represent the total number of metabolites that participate in a reaction (Fig. 1B).

Though widely studied [8, 16], the RAG has known limitations and overlooks key aspects of the connectivity of metabolic networks. The RAG does not distinguish between forward and backward fluxes, nor does it incorporate the irreversibility of reactions (by construction **A** is a symmetric matrix). Furthermore, the structure of **A** is dominated by the large number of edges introduced by pool metabolites that appear in many reactions, such as water, ions or enzymatic cofactors. Computational schemes have been introduced to mitigate the bias caused by pool metabolites [27], but these do not follow from biophysical considerations and need manual calibration. Finally, the construction of the graph **A** from is not easily extended to incorporate the effect of environmental changes.

### B. Metabolic graphs that incorporate flux directionality and biological context

To address the limitations of the reaction adjacency graph **A**, we propose a graph formulation that follows from a flux-based perspective. To construct our graph, we unfold each reaction into two separate directions (forward and reverse) and redefine the links between reaction nodes to reflect producer-consumer relationships. Specifically, two reactions are connected if one produces a metabolite that is consumed by the other. As shown below, this definition leads to graphs that naturally account for the reversibility of reactions, and allows for the seamless integration of biological contexts modelled through FBA.

Inspired by matrix formulations of chemical reaction network kinetics [36], we rewrite the reaction rate vector **v** as:

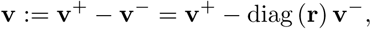

where **v**^+^ and **v−** are non-negative vectors containing the forward and backward reaction rates, respectively. Here the *m × m* matrix diag (**r**) contains **r** in its main diagonal, and **r** is the *m*-dimensional reversibility vector with components *r_j_* = 1 if reaction *R*_*j*_ is reversible and *r*_*j*_ = 0 if it is irreversible. With these definitions, we can rewrite the metabolic model in (Eq. 2) as:

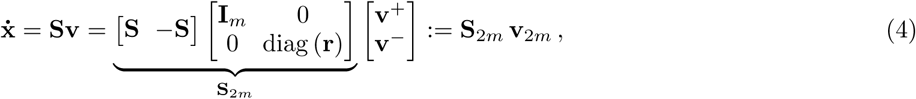

where **v**_2*m*_ := [**v**^+^ **v−**]^*T*^ is the unfolded 2*m*-dimensional vector of reaction rates, **I**_*m*_ is the *m × m* identity matrix, and we have defined **S**_2*m*_, the unfolded version of the stoichiometric matrix of the 2*m* forward and reverse reactions.

#### 1. Probabilistic Flux Graph: a directional blueprint of metabolism

The unfolding into forward and backward fluxes leads us to the definition of *production* and *consumption* stoichio-metric matrices:

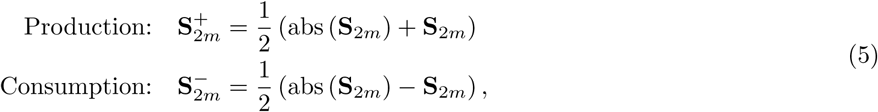

where abs (**S**_2*m*_) is the matrix of absolute values of the corresponding entries of **S**_2*m*_. Note that each entry of the matrix 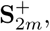, denoted 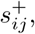, gives the number of molecules of metabolite *X*_*i*_ produced by reaction *R*_*j*_. Conversely, the entries of 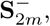, denoted 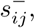, correspond to the number of molecules of metabolite *X*_*i*_ consumed by reaction *R*_*j*_. Within our directional flux framework, it is natural to consider a purely probabilistic description of producer-consumer relationships between reactions, as follows. Suppose we are given a stoichiometric matrix **S** without any additional biological information, such as metabolite concentrations, reaction fluxes, or kinetic rates. In the absence of such information, the probability that metabolite *X_k_* is produced by reaction *R_i_* and consumed by reaction *R_j_* is:

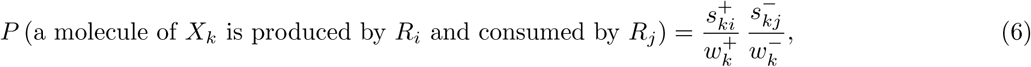

where 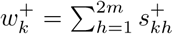 and 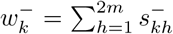 are the total number of molecules of *X_k_* produced and consumed by all reactions. Unlike models in that rely on stochastic chemical kinetics [37], the probabilities in (Eq. 6) do not contain information on kinetic rate constants, which are typically not available for genome-scale metabolic models [38]. In our formulation, the relevant probabilities contain only the stoichiometric information included in the matrix **S**_2*m*_ and should not be confused with the reaction propensity functions in Gillespie-type stochastic simulations of biochemical systems.

We thus define the weight of the edge between reaction nodes *R_i_* and *R_j_* as the probability that *any* metabolite chosen at random is produced by *R_i_* and consumed by *R_j_*. Summing over all metabolites and normalizing, we obtain the edge weights of the adjacency matrix of the PFG:

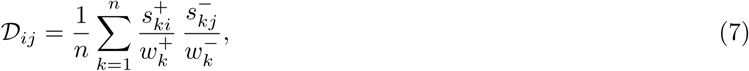

in which 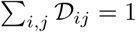 (i.e., the probability that any metabolite is consumed/produced by any reaction is 1). Rewritten compactly in matrix form, we obtain the

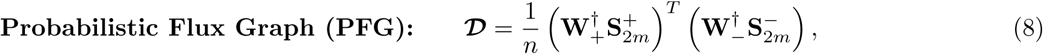

where 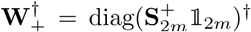, 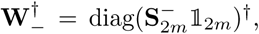 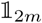 is a vector of ones, and † denotes the Moore-Penrose pseudoinverse. In Figure 1C we illustrate the creation of the PFG for a toy network. The PFG is a weighted, directed graph which encodes a blueprint of the whole metabolic model, and provides a natural scaling of the contribution of pool metabolites to flux transfer. We remark that the PFG is distinct from directed analogues of the RAG constructed from boolean production and consumption stoichiometric matrices, as shown in Sec. SI 1.

We now extend the construction of the PFG to accommodate specific environmental contexts or growth conditions.

Cells adjust their metabolic fluxes to respond to the availability of nutrients and environmental requirements. Flux Balance Analysis (FBA) is a widely used method to predict environment-specific flux distributions. FBA computes a vector of metabolic fluxes **v∗** that maximise a cellular objective (e.g., biomass, growth or ATP production). The FBA solution is obtained assuming steady state conditions (**ẋ** = 0 in (Eq. 2)) subject to constraints that describe the availability of nutrients and other extracellular compounds [16]. The core elements of FBA are briefly summarised in Section A 1.

To incorporate the biological information afforded by FBA solutions into the structure of a metabolic graph, we again define the graph edges in terms of production and consumptions fluxes. Similarly to (Eq. 4), we unfold the FBA solution vector **v\*** into forward and backward components: positive entries in the FBA solution correspond to forward fluxes, negative entries in the FBA solution correspond to backward fluxes. From the unfolded fluxes

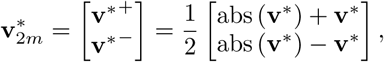

we compute the vector of production and consumption fluxes as

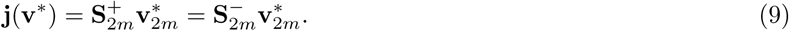

The *k*-th entry of 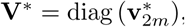 is the flux at which metabolite *X*_*k*_ is produced and consumed, and the equality of the production and consumption fluxes follows from the steady state condition, **x˙** = 0.

To construct the flux graph, we define the weight of the edge between reactions *R_i_* and *R_j_* as the *total flux of metabolites produced by R_i_ that are consumed by R_j_*. Assuming that the amount of metabolite produced by one reaction is distributed among the reactions that consume it in proportion to their flux (and respecting the stoichiometry), the flux of metabolite *X_k_* from reaction *R_i_* to *R_j_* is given by

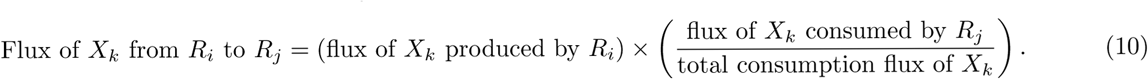

For example, if the total flux of metabolite *X_k_* is 10 mmol*/*gDW*/*h, with reaction *R_i_* producing *X_k_* at a rate 1.5 mmol/gDW/h and reaction *R_j_* consuming *X_k_* at a rate 3.0 mmol/gDW/h, then the flux of *X_k_* from *R_i_* to *R_j_* is 0.45 mmol/gDW/h.

Summing (10) over all metabolites, we obtain the edge weight relating reactions *R_i_* and *R_j_*:

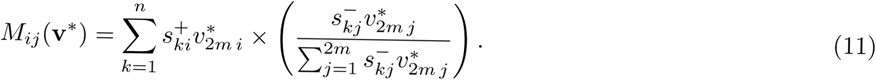

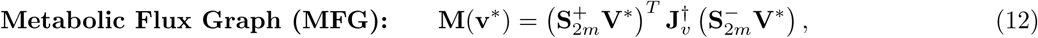

where 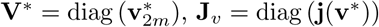 and † denotes the matrix pseudoinverse. The MFG is a directed, weighted graph with edge weights in units of mmol/gDW/h. Self-loops describe the metabolic flux of autocatalytic reactions, i.e., those in which products are also reactants.

The MFG provides a versatile framework to create environment-specific metabolic graphs from FBA solutions. In Figure 1C, we illustrate the creation of MFGs for a toy network under different biological scenarios. In each case, an FBA solution is computed under a fixed uptake flux with the remaining fluxes constrained to account for differences in the biological environment: in scenario 1, the fluxes are constrained to be strictly positive and no larger than the nutrient uptake flux, while in scenario 2 we impose a positive lower bound on reaction *R*_7_. Note how the MFG for scenario 2 displays an extra edge between reactions *R*_4_ and *R*_7_, as well as distinct edge weights to scenario 1 (see Sec. SI 2 for details). These differences illustrate how changes in the FBA solutions translate into different graph connectivities and edge weights.

### C. Flux-based graphs of *Escherichia coli* metabolism

To illustrate our framework, we construct and analyse the flux graphs of the well-studied core metabolic model of *coli* [33]. This model (Fig. 2A) contains 72 metabolites and 95 reactions, grouped into 11 pathways, which describe the main biochemical routes in central carbon metabolism [40–42]. We provide a Supplemental Spreadsheet with full details of the reactions and metabolites in this model, as well as all the results presented below.

**FIG. 2.**
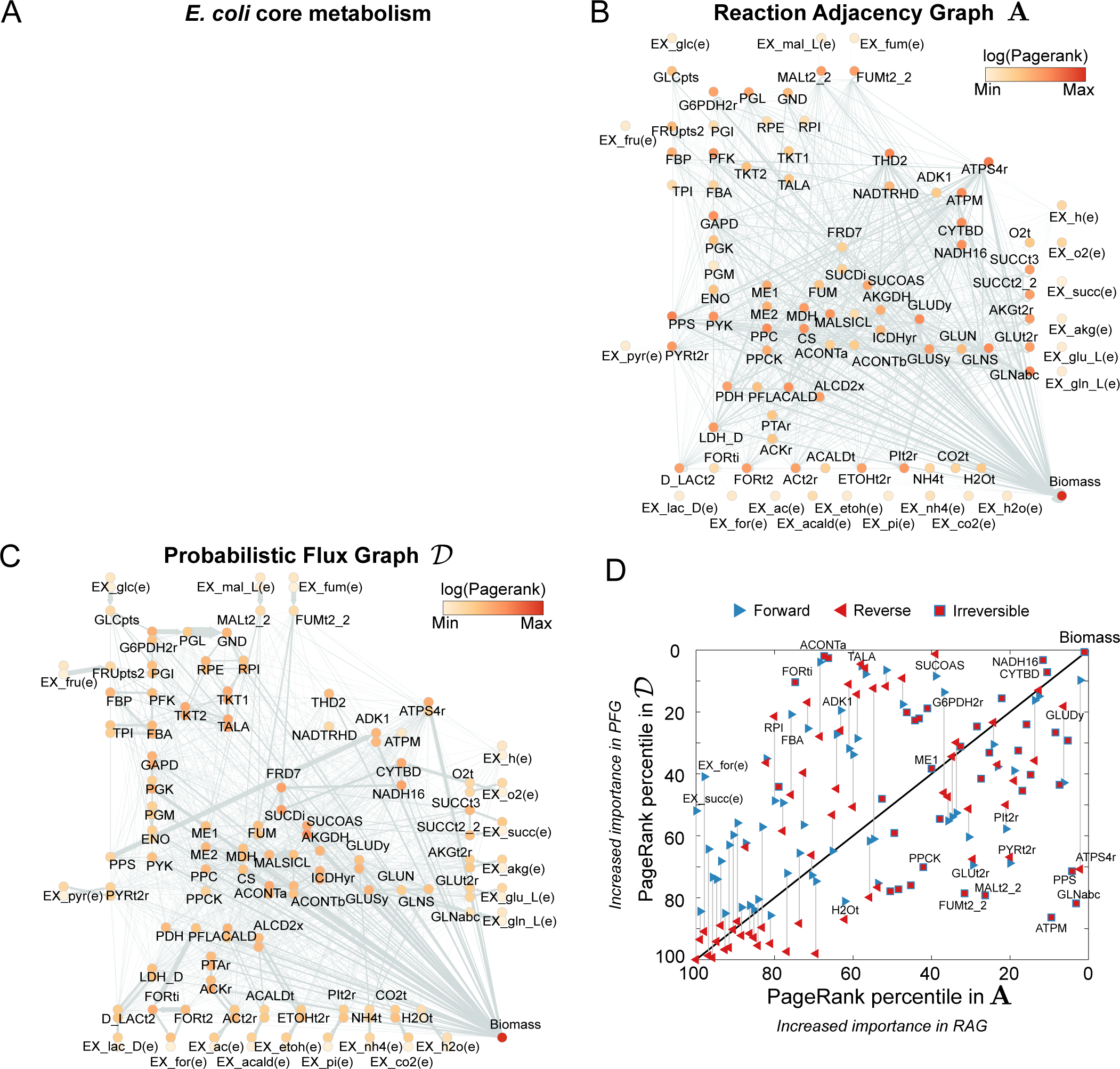
Graphs for the core metabolism of *Escherichia coli*. (A) Map of the *E. coli* core metabolic model created with the online tool Escher [33, 39]. (B) The standard Reaction Adjacency Graph **A**, as given by (Eq. 3). The nodes represent reactions; two reactions are linked by an undirected edge if they share reactants or products. The nodes are coloured according to their PageRank score, a measure of their centrality (or importance) in the graph. (C) The directed Probabilistic Flux Graph **𝒟**, as computed from (Eq. 8). The reversible reactions are unfolded into two overlapping nodes (one for the forward reaction, one for the backward). The directed links indicate flow of metabolites produced by the source node and consumed by the target node. The nodes are coloured according to their PageRank score. (D) Comparison of PageRank percentiles of reactions in **A** and **𝒟**. Reversible reactions are represented by two triangles connected by a line; both share the same PageRank in **A**, but each has its own PageRank in **𝒟**. Reactions that appear above (below) the diagonal have increased (decreased) PageRank in **𝒟** as compared to **A**.

### 1. The Probabilistic Flux Graph: the impact of directionality

To examine the effect of flux directionality on the metabolic graphs, we compare the Reaction Adjacency Graph (**A**) and our proposed Probabilistic Flux Graph (**𝒟**) for the same metabolic model in Figure 2. The **A** graph has 95 nodes and 1,158 undirected edges, whereas the **𝒟** graph has 154 nodes and 1,604 directed and weighted edges. The increase in node count is due to the unfolding of forward and backward reactions into separate nodes. Unlike the **A** graph, where edges represent shared metabolites between two reactions, the directed edges of the **𝒟** graph represent the flow of metabolites from a source to a target reaction. A salient feature of both graphs is their high connectivity, which is not apparent from the traditional pathway representation in Figure 2A.

The effect of directionality becomes apparent when comparing the importance of reaction nodes in both graphs (Figure 2B–D), as measured with the PageRank score for node centrality [43, 44]. The overall node hierarchy is maintained across both graphs: exchange reactions tend to have low PageRank centrality scores, core metabolic reactions have high scores, and the biomass reaction has the highest scores in both graphs. Yet we also observe substantial changes in specific reactions. For example, the reactions for ATP maintenance (ATPM, irreversible), phosphoenolpyruvate synthase (PPS, irreversible) and ABC-mediated transport of L-glutamine (GLNabc, irreversible) drop from being among the top 10% most important reactions in the **A** graph to the bottom percentiles in the **𝒟** graph. Conversely, reactions such as aconitase A (ACONTa, irreversible), transaldolase (TALA, reversible) and succinyl-CoA synthetase (SUCOAS, reversible), and formate transport via diffusion (FORti, irreversible) gain substantial importance in the **𝒟** graph. For instance, FORti is the sole consumer of formate, which is produced by pyruvate formate lyase (PFL), a reaction that is highly connected to the rest of the network. Importantly, in most of the reversible reactions, such as ATP synthase (ATPS4r), there is a wide gap between the PageRank of the forward and backward reactions, suggesting a marked asymmetry in the importance of metabolic flows.

Community detection is frequently used in the analysis of complex graphs: nodes are clustered into tightly related communities that reveal the coarse-grained structure of the graph, potentially at different levels of resolution [47–49]. The community structure of metabolic graphs has been the subject of multiple analyses [12, 14, 47]. However, most community detection methods are applicable to undirected graphs only, and thus fail to capture the directionality of the metabolic graphs we propose here. To account for graph directionality, we use the Markov Stability community detection framework [49–51], which uses diffusion on graphs to detect groups of nodes where flows are retained persistently across time scales. Markov Stability is ideally suited to find multi-resolution community structure [48] and can deal with both directed and undirected graphs [49, 52] (see Sec. A 2). In the case of metabolic graphs, Markov Stability can reveal groups of reactions that are closely interlinked by the flow of metabolites that they produce and consume.

Figure 3 shows the difference between the community structure of the undirected RAG and the directed PFG of the core metabolism of *E. coli*. For the **A** graph, Markov Stability reveals a partition into seven communities (Figure 3B, see also Sec. SI 3), which are largely dictated by the many edges created by shared pool metabolites. For example, community C1(**A**) is mainly composed of reactions that consume or produce ATP and water. Yet, the biomass reaction (the largest consumer of ATP) is not a member of C1(**A**) because, in the standard **A** graph construction, any connection involving ATP has equal weight. Other communities in **A** are also determined by pool metabolites, e.g. C2(**A**) is dominated by H^+^, and C3(**A**) is dominated by NAD^+^ and NADP^+^, as illustrated by word clouds of the relative frequency of metabolites in the reactions within each community. The community structure in **A** thus reflects the limitations of the RAG construction due to the absence of biological context and the large number of uninformative links introduced by pool metabolites.

**FIG. 3.**
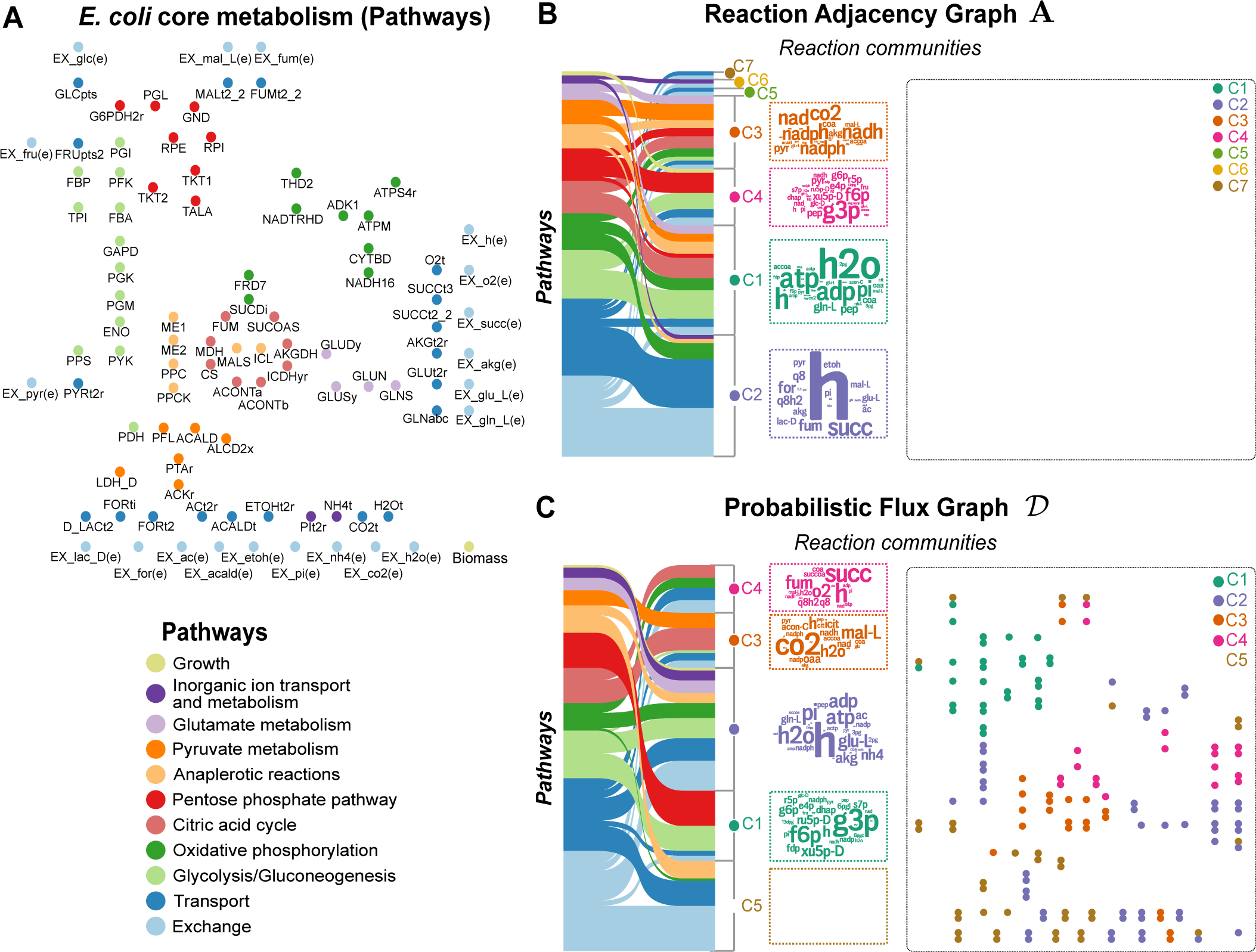
Directionality and community structure of graphs for *Escherichia coli* metabolism. (A) Reactions of the core model of *E. coli* metabolism grouped into eleven biochemical pathways, (B–C) Graphs **A** and **𝒟** from Fig. 2B–C partitioned into communities computed with the Markov Stability method; for clarity, the graph edges are not shown. The Sankey diagrams [45, 46] show the correspondence between biochemical pathways and the communities found in each graph. The word clouds contain the metabolites that participate in the reactions each community, and the word size is proportional to the number of reactions in which each metabolite participates.

For the **𝒟** graph, we found a robust partition into five communities (Figure 3C, see also Sec. SI 3), which comprise reactions related consistently through biochemical pathways. Community C1(**𝒟**) contains the reactions in the pentose phosphate pathway together with the first steps of glycolysis involving D-fructose, D-glucose, or D-ribulose. Community C2(**𝒟**) contains the main reactions that produce ATP from substrate level as well as oxidative phosphorylation and the biomass reaction. Community C3(**𝒟**) includes the core of the citric acid cycle, anaplerotic reactions related to malate syntheses, as well as the intake of cofactors such as CO_2_. Community C4(**𝒟**) contains reactions that are secondary sources of carbon (such as malate and succinate), as well as oxidative phosphorilation reactions. Finally, community C5(**𝒟**) contains reactions that are part of the pyruvate metabolism subsystem, as well as transport reactions for the most common secondary carbon metabolites such as lactate, formate, acetaldehyde and ethanol. Altogether, the communities of the **𝒟** graph reflect metabolite flows associated with specific cellular functions, as a consequence of including flux directionality in the graph construction. As seen in Fig. 3C, the communities are no longer exclusively determined by pool metabolites (e.g., water is no longer dominant and protons are spread among all communities). For a more detailed explanation and comparison of the communities found in the **A** and **𝒟** and graphs, see Section SI 3. Full information about PageRank scores and communities is provided in the Supplementary Spreadsheet.

### 2. Metabolic Flux Graphs: the impact of growth conditions and biological context

To incorporate the impact of environmental context, we construct Metabolic Flux Graphs in (Eq. 12) using FBA solutions of the core model of *E. coli* metabolism in four relevant growth conditions: aerobic growth in rich media with glucose; aerobic growth in rich media with ethanol, anaerobic growth in glucose; and aerobic growth in glucose but phosphate- and ammonium-limited. The results in Figure 4 show how changes in metabolite fluxes under different biological contexts have a direct effect in the MFG. Note that, in all cases, the MFGs have fewer nodes than the blueprint graph **𝒟** since the FBA solutions contain numerous reactions with zero flux.

**FIG. 4.**
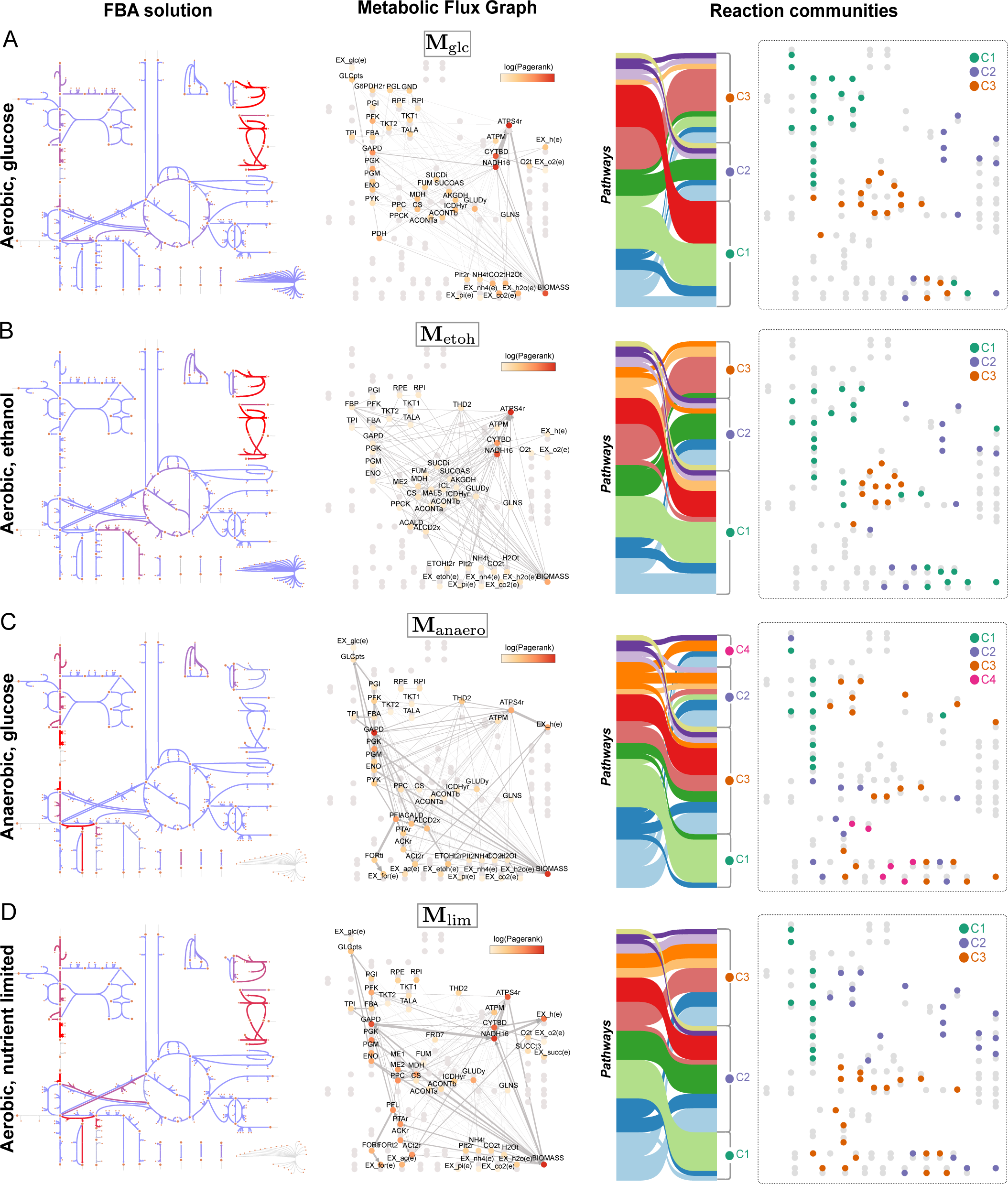
Metabolic Flux Graphs for *Escherichia coli* in different growth conditions. The MFGs are computed from (Eq. 12) and the FBA solutions in four different environments: (A) aerobic growth in D-glucose, (B) aerobic growth in ethanol, (C) anaerobic growth in D-glucose, and (D) aerobic growth in D-glucose but with limited ammonium and phosphate. Each subfigure shows: (left) flux map obtained with Escher [39], where the increased red colour of the arrows indicates increased flux; (centre) Metabolic Flux Graph with nodes coloured according to their PageRank (zero flux reactions are in grey; thickness of connections proportional to fluxes); (right) community structure computed with the Markov Stability method together with Sankey diagrams showing the correspondence between biochemical pathways and MFG communities.

Next we summarise how the changes in the community structure of the MFGs for the four conditions reflect the distinct relationships of functional pathways in response to growth requirements.

*a. Aerobic growth in D-glucose (***M**_glc_). We found a robust partition into three communities with an intuitive biological interpretation (Fig. 4A and Fig. SI 2A). C1(**M**_glc_) is the carbon-processing community, comprising reactions that process carbon from D-glucose to pyruvate including most of the glycolysis and pentose phosphate pathways, together with related transport and exchange reactions. C2(**M**_glc_) harbours the bulk of reactions related to oxidative phosphorylation and the production of energy in the cell, including the electron transport chain of NADH dehydro-genase, cytochrome oxidase, and ATP synthase, as well as transport reactions for phosphate and oxygen intake and proton balance. C2(**M**_glc_) also includes the growth reaction, consistent with ATP being the main substrate for both the ATP maintenance (ATPM) requirement and the biomass reaction in this growth condition. Finally, C3(**M**_glc_) contains reactions related to the citric acid cycle (TCA) and the production of NADH and NADPH (i.e., the cell’s reductive power), together with carbon intake routes strongly linked to the TCA cycle, such as those starting from phosphoenolpyruvic acid (PEP).

*b. Aerobic growth in ethanol (***M**_etoh_). The robust partition into three communities that we found for this scenario resembles the structure of **M**_glc_ with subtle, yet important, differences (Fig. 4B and Fig. SI 2B). Most salient are the differences in the carbon-processing community C1(**M**_etoh_), which reflects the switch from D-glucose to ethanol as a carbon source. C1(**M**_etoh_) contains gluconeogenic reactions (instead of glycolytic), due to the reversal of flux induced by the change of carbon source, as well as anaplerotic reactions and reactions related to glutamate metabolism. In particular, the reactions in this community are related to the production of precursors such as PEP, pyruvate, 3-phospho-D-glycerate (3PG), glyceraldehyde-3-phosphate (G3P), D-fructose-6-phosphate (F6P), and D-glucose-6- phosphate, all of which are substrates for growth. Consequently, the biomass reaction is also grouped within C1(**M**_etoh_) due to the increased metabolic flux of precursors relative to ATP production in this biological scenario. The other two reaction communities (energy-generation C2(**M**_etoh_) and citric acid cycle C3(**M**_etoh_)) display less prominent differences relative to the **M**_glc_ graph, with additional pyruvate metabolism and anaplerotic reactions as well as subtle ascriptions of reactions involved in NADH/NADPH balance and the source for acetyl-CoA.

*c. Anaerobic growth in D-glucose (***M**_anaero_). The profound impact of the absence of oxygen on the metabolic balance of the cell is reflected in drastic changes in the MFG (Fig. 4C and Fig. SI 2C). Both the connectivity and reaction communities in the MFG are starkly different from the aerobic scenarios, with a much diminished presence of oxidative phosphorylation and the absence of the first two steps of the electron transport chain (CYTBD and NADH16). We found that **M**_anaero_ has a robust partition into four communities. C1(**M**_anaero_) still contains carbon processing (glucose intake and glycolysis), yet now decoupled from the pentose phosphate pathway. C3(**M**_anaero_) in-cludes the pentose phosphate pathway grouped with the citric acid cycle (incomplete) and the biomass reaction, as well as the growth precursors including alpha-D-ribose-5-phosphate (r5p), D-erythrose-4-phosphate (e4p), 2-oxalacetate and NADPH. The other two communities are specific to the anaerobic context: C2(**M**_anaero_) contains the conversion of PEP into formate (more than half of the carbon secreted by the cell becomes formate [53]); C4(**M**_anaero_) includes NADH production and consumption via reactions linked to glyceraldehyde-3-phosphate dehydrogenase (GAPD).

*d. Aerobic growth in D-glucose but limited phosphate and ammonium (***M**_lim_). Under growth-limiting conditions, we found a robust partition into three communities (Fig. 4D and Fig. SI 2D). The community structure reflects *overflow metabolism* [54], which occurs when the cell takes in more carbon than it can process. As a consequence, the excess carbon is secreted from the cell, leading to a decrease in growth and a partial shutdown of the citric acid cycle.

This is reflected in the reduced weight of the TCA pathway and its grouping with the secretion routes of acetate and formate within C3(**M**_lim_). Hence, C3(**M**_lim_) comprises reactions that are not strongly coupled in favourable growth conditions, yet are linked together by metabolic responses to limited ammonium and phosphate. Furthermore, the carbon-processing community C1(**M**_lim_) contains the glycolytic pathway, yet detached from the pentose phosphate pathway (as in **M**_anaero_), highlighting its role in precursor formation. The bioenergetic machinery, contained in community C2(**M**_lim_), includes the pentose phosphate pathway, with a smaller role for the electron transport chain (21.8% of the total ATP as compared to 66.5% in **M**_glc_).

In addition to the effect on community structure, Figure 4 also shows the changes induced by the environment on the MFG connectivity and relative importance of reactions, as measured by their PageRank score. To provide a global snapshot of the effect of growth conditions on cellular metabolism, Figure 5 shows the cumulative PageRank of each pathway for each of the MFGs. The cumulative PageRank quantifies the relative importance of pathways, and how their importance changes upon environmental shifts. In aerobic growth, a shift from glucose to ethanol (**M**_glc_ → **M**_etoh_) as carbon source increases the importance of pyruvate metabolism and oxidative phosphorylation, while reducing the importance of the pentose phosphate pathway. A shift from aerobic to anaerobic growth in glucose (**M**_glc_ → **M**_anaero_) sees a large reduction in the importance of oxidative phosphorylation and the citric acid cycle, coupled with a large increase in the importance of gluconeogenesis, pyruvate metabolism, and transport and exchange reactions. The effect of growth-limiting conditions in aerobic growth under glucose (**M**_glc_ → **M**_lim_) is reflected on the increased importance of pyruvate metabolism and a reduction in the importance of oxidative phosphorylation, citric acid cycle, and the pentose phosphate pathway. The importance of transport and exchange reactions is also increased under limiting conditions. Such qualitative relations between growth conditions and the importance of specific pathways highlights the utility of the MFGs to characterise systemic metabolic changes in response to environmental conditions.

**FIG. 5.**
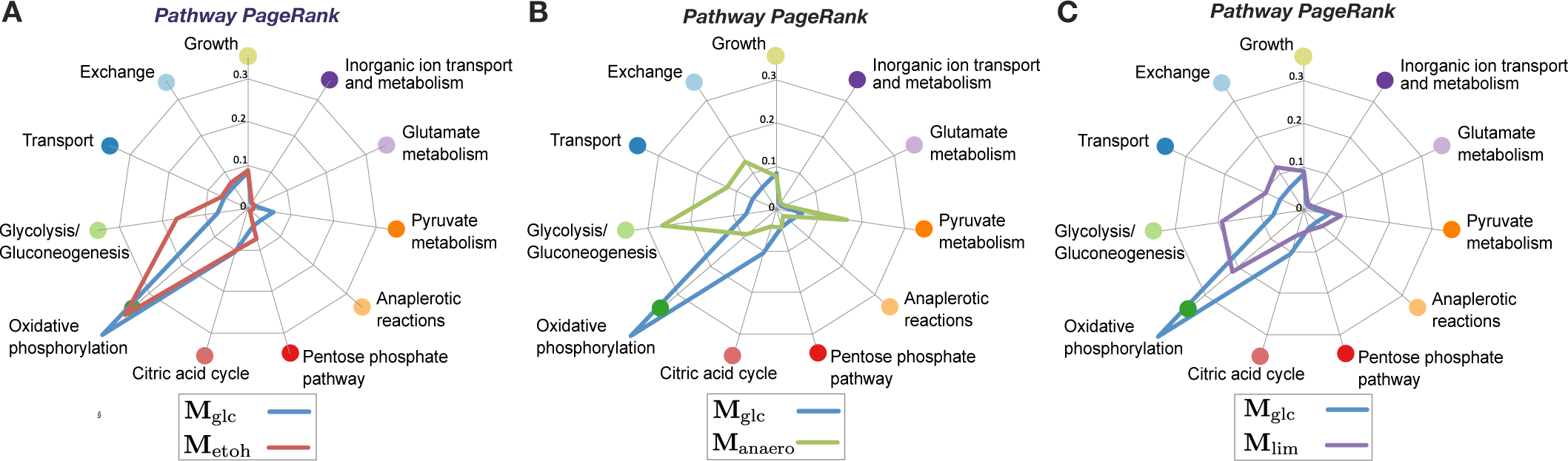
Pathway centrality (PageRank) computed from the MFG of different growth conditions. The cumulative pathway PageRank reflects the relative importance of metabolic pathways in each MFG. Changes in pathway centrality indicate the overall rearrangement of fluxes within the pathways in response to environmental shifts: (A) from aerobic glucose-rich to aerobic ethanol-rich; (B) from aerobic glucose-rich to anerobic glucose-rich; (C) from aerobic glucose-rich to a similar medium with limited phosphate and ammonium. Variations in cumulative Pagerank highlight changes across most cellular pathways.

A more detailed discussion of the changes in pathways and reactions can be found in Section SI 4 and Fig. SI 2 in the Supplementary Information, with full details of all the results in the Supplemental Spreadsheet.

## 3. Multiscale organisation of metabolic flux graphs

The definition of the MFGs as *directed graphs* opens up the application of network-theoretic tools for detecting modules of reaction nodes and the hierarchical relationships among them. In contrast with methods for undirected graphs, the Markov Stability framework [50, 55] can be used to detect multi-resolution community structure in directed graphs (Sec. A 2), thus allowing the exploration of the multiscale organisation of metabolic reaction networks. The modules so detected reflect subsets of reactions where metabolic fluxes tend to be contained.

Figure 6 illustrates this multiscale analysis on **M**_glc_, the MFG of *E. coli* under aerobic growth in glucose. By varying the Markov time *t*, a parameter in the Markov Stability method, we scanned the community structures at different resolutions. Our results show that, from finer to coarser resolution, the MFG can be partitioned into 11, 7, 5, 3, and 2 communities of high persistence across Markov time (extended plateaux over *t*, as shown by low values of *V I*(*t, t^!^*)) and high robustness under optimisation (as shown by dips in *V I*(*t*)). For further details, see Section A 2 and Refs. [48–50, 55].

**FIG. 6.**
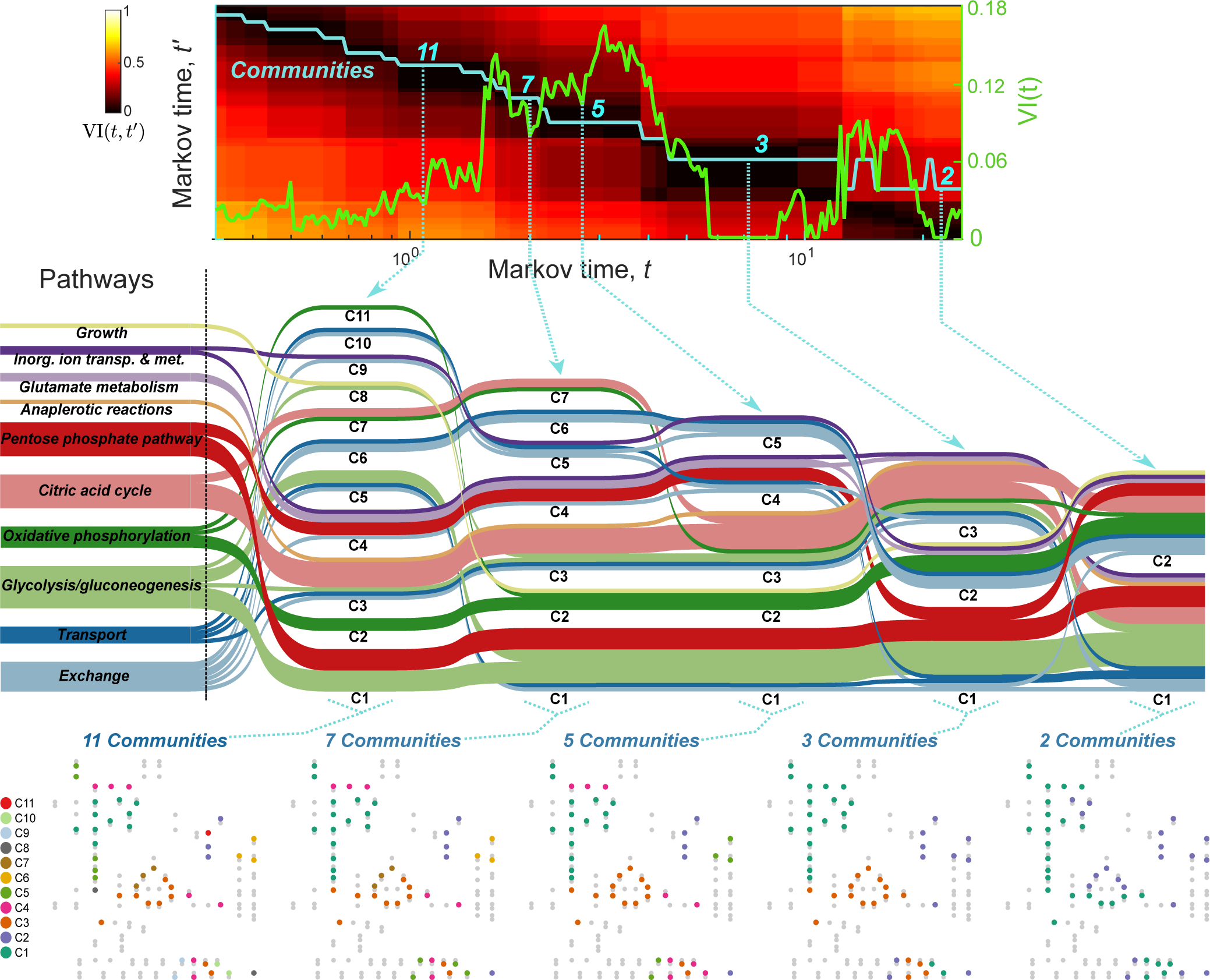
Community structure of flux graphs across different scales. We applied the Markov Stability method to partition the metabolic flux graph for *E. coli* aerobic growth in glucose (**M**_glc_) across levels of resolution. The top panel shows the number of communities of the optimal partition (blue line) and two measures of its robustness (*V I*(*t*) (green line) and *V I*(*t, t^!^*) (colour map)) as a function of the Markov time *t* (see text and Methods section). The five Markov times selected correspond to robust partitions of the graph into 11, 7, 5, 3, and 2 communities, as signalled by extended low values of *V I*(*t, t^!^*) and low values (or pronounced dips) of *V I*(*t*). The Sankey diagram (middle panel) visualises the multiscale organisation of the communities of the flux graph across Markov times, and the relationship of the communities with the biochemical pathways. The bottom panel shows the five partitions at the selected Markov times. The partition into 3 communities corresponds to that in Figure 4A.

The Sankey diagram in Fig. 6 visualises the pathway composition of the graph partitions and their relationships across different resolutions. As we decrease the resolution (longer Markov times), the reactions in different pathways assemble and split into different groupings, reflecting both specific relationships and general organisation principles associated with this growth condition. A general observation is that glycolysis is grouped together with oxidative phosphorylation across most scales, underlining the fact that those two pathways function as cohesive metabolic sub-units in aerobic conditions. In contrast, the exchange and transport pathways appear spread among multiple partitions across all resolutions. This is expected, as exchange/transport are enabling functional pathways, in which reactions do not interact amongst themselves but rather feed substrates to other pathways.

Other reaction groupings reflect more specific relationships. For example, the citric acid cycle (always linked to anaplerotic reactions) appears as a cohesive unit across most scales, and only splits in two in the final grouping, reflecting the global role of the TCA cycle in linking to both glycolysis and oxidative phosphorylation. The pentose phosphate pathway, on the other hand, is split into two groups (one linked to glutamate metabolism and another one linked to glycolysis) across early scales, only merging into the same community towards the final groupings. This suggests a more interconnected flux relationship of the different steps of the penthose phosphate pathway with the rest of metabolism. In Figure SI 2, we present the multiscale analyses of the reaction communities for the other three growth scenarios (**M**_etoh_, **M**_anaero_, **M**_lim_).

### D. Using MFGs to analyse hepatocyte metabolism in wild type and PH1 mutant human cells

To showcase the applicability of our framework to larger metabolic models, we analyse a model of human hepatocyte (liver) metabolism with 777 metabolites and 2589 reactions [34], which extends the widely used HepatoNet1 model [56] with an additional 50 reactions and 8 metabolites. This extended model was used in Ref. [34] to compare wild type cells (WT) and cells affected by the rare disease Primary Hyperoxaluria Type 1 (PH1), which lack alanine:glyoxylate aminotransferase (AGT) due to a genetic mutation. AGT is an enzyme found in peroxisomes and its mutation decreases the breakdown of glyoxylate, with subsequent accumulation of calcium oxalate that leads to liver damage.

Following [34], we first obtain 442 FBA solutions for different sets of metabolic objectives for both the wild type (WT) model and the PH1 model lacking AGT (reaction r2541). We then generate the corresponding 442 MFGs for each WT and PH1, and obtain the averages over each ensemble: 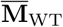 and 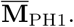. Of the 2589 reactions in the model, 2448 forward and 1362 reverse reactions are present in at least one of the FBA solutions. Hence the average MFGs have 3810 nodes each (see Supplementary Spreadsheet for full details about the reactions).

Figure 7A shows the MFG for the wild type 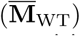 coloured according to a robust partition into 7 communities obtained with Markov Stability. The seven communities are broadly linked to amino acid metabolism (C0), energy metabolism (C1 and C5), glutathione metabolism (C2), fatty acid and bile acid metabolism (C3 and C4) and choles-terol metabolism and lipoprotein particle assembly (C6). As expected, the network community structure of the MFG is largely preserved under the AGT mutation: the Sankey diagramme in Fig. 7B shows a remarkable match between the partitions of 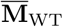 and 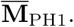 found independently with Markov Stability. Despite this similarity, our method also identified subtle but important differences between the healthy and diseased networks. In particular, C3’ in 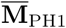 receives 60 reactions, almost all taking place in the peroxisome and linked to mevalonate and iso-pentenyl pathways, as well as highly central transfer reactions of PP_*i*_, O_2_ and H_2_O_2_ between the peroxisome and the cytosol (r1152, r0857, r2577).

**FIG. 7.**
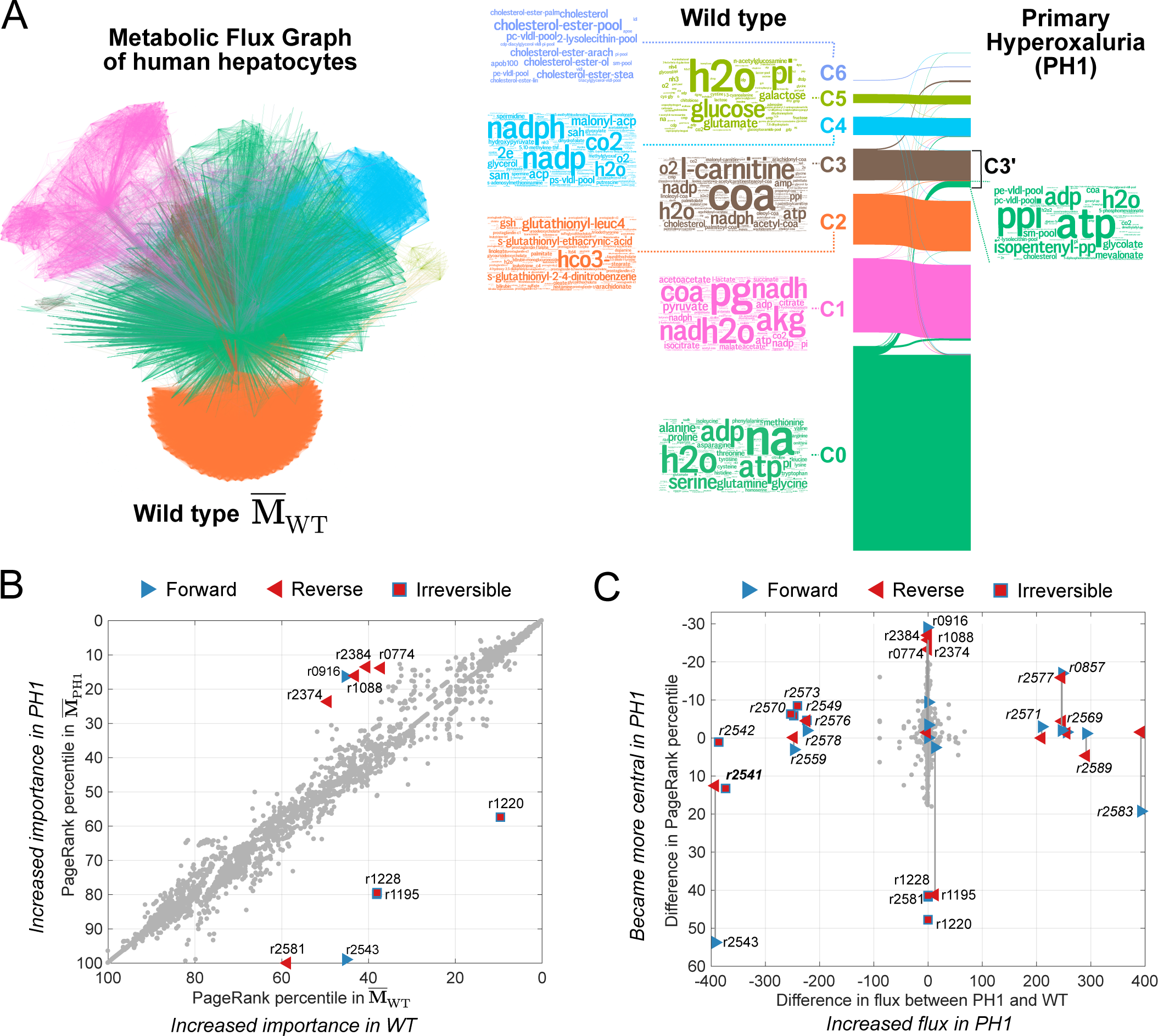
MFG analysis of a model of human hepatocyte metabolism and the genetic condition PH1. Average MFG of wild-type hepatocytes cells over 442 metabolic objectives. The reaction nodes are coloured according to communities in a 7-way partition obtained with Markov Stability. The Sankey diagramme shows the consistency between the communities in the wild type MFG and the communities independently found in the MFG of the mutated PH1 cells. Word clouds of the most frequent metabolites in the reactions of the WT communities reveal functional groupings (see text). Under the PH1 mutation, the only large change relates to metabolites that join C3’ from community C0 in WT. (B) Comparison of the PageRank percentiles in the WT and PH1 MFGs, with reactions whose rank changes by more than 20 percentiles labelled. (C) Difference in FBA flux between WT and PH1 vs difference in PageRank percentile between WT and PH1. Reactions whose flux difference is greater than 100mmol/gDW/h (italics) or whose change in PageRank percentile is greater than 20 are labelled. The differences in centrality (PageRank) provide complementary information, revealing additional important reactions affected by the PH1 mutation that knocks out reaction r2541.

Overall, the centrality (PageRank) of most reactions in the MFG is relatively unaffected by the PH1 mutation, as shown by the good correlation between the PageRank percentiles in 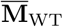 and 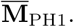 in Figure 7C. Yet, there are notable exceptions, and the reactions that exhibit the largest change in PageRank centrality (labelled in Fig. 7B) provide biological insights into the disease state. Specifically, the four reactions (r0916, r1088, r2384, r2374) that undergo the largest increase in centrality from 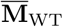 to 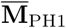 are related to the transfer of citrate out of the cytosol in exchange for oxalate and PEP; whereas those with the largest decrease of PageRank from 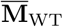 to 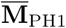 are related to VLDL-pool reactions (r1228, r1195, r1220) and to transfers of hydroxypyruvate and alanine from peroxisome to cytosol (r2581, r2543). It is worth remarking that although oxalate and citrate reactions are directly linked to metabolic changes associated with the PH1 diseased state, none of them exhibits large changes in their flux predicted by FBA, yet they show large changes in PageRank centrality.

These observations underscore how the information provided by our network analysis provides complementary information to the analysis of FBA fluxes alone. As shown in Figure 7D, there is a group of reactions (labelled with italics in the Figure) that exhibit large gains or decreases in their flux under the PH1 mutation, yet they only undergo relatively small changes in their PageRank scores. Closer inspection reveals that most of these reactions are close to the AGT reaction (r2541, highlighted in the Figure) in the pathway and involve the conversion of glycolate, pyruvate, glycine, alanine and serine. Hence the changes in flux follow from the *local rearrangement* of flows as a consequence of the deletion of reaction r2541. On the other hand, the citrate and oxalate reactions (r0916, r1088, r2384, r2374) with large changes in their centrality yet undergo small changes in flux, thus reflecting *global changes* in the flux structure of the network. Importantly, the transport reactions of O_2_, H_2_O_2_, serine and hydroxypyruvate between cytosol and peroxisome (r0857, r2577, r2583, r2543) all undergo large changes both in centrality and flux, highlighting the importance of peroxisome transfer reactions in PH1. We provide a full spreadsheet with these analyses as Supplementary Material for the interested reader.

## DISCUSSION

Metabolism is commonly understood in terms of functional pathways interconnected into metabolic networks [39], i.e., metabolites linked by arrows representing enzymatic reactions between them as in Figure 2A. However, such standard representations are not amenable to rigorous graph-theoretic analysis. Fundamentally different graphs can be constructed from such metabolic reactions depending on the chosen representation of species/interactions as nodes/edges, e.g., reactions as nodes; metabolites as nodes; or both reaction and metabolites as nodes [16]. Each of those graphs can be directed or undirected and with weighted links computed according to different rules. The choices and subtleties in graph construction are crucial both to capture the relevant metabolic information and to interpret their topological properties [10, 17].

Here, we have presented a flux-based strategy to build graphs for metabolic networks. Our graphs have reactions as nodes and directed weighted edges representing the flux of metabolites produced by a source reaction and consumed by a target reaction. This principle is applied to build both ‘blueprint’ graphs (PFG), which summarise probabilistically the fluxes of the whole metabolism of an organism, as well as context-specific graphs (MFGs), which reflect specific environmental conditions. The blueprint Probabilistic Flux Graph has edge weights equal to the probability that source/target reactions produce/consume a molecule of a metabolite chosen at random in the absence of any other information, and can thus be used when the stoichiometric matrix is the only information available. The PFG construction naturally tames the over-representation of pool metabolites without the need to remove them from the graph arbitrarily, as often done in the literature [26, 28–30]. Context-specific Metabolic Flux Graphs (MFGs) can incorporate the effect of the environment, e.g., with edge weights corresponding to the total flux of metabolites between reactions as computed by Flux Balance Analysis (FBA). FBA solutions for different environments can then be used to build different metabolic graphs in different growth conditions.

The proposed graph constructions provide complementary tools for studying the organisation of metabolism and can be embedded into any FBA-based modelling pipeline. Specifically, the PFG relies on the availability of a well-curated stoichiometric matrix, which is produced with metabolic reconstruction techniques that typically precede the application of FBA. The MFG, on the other hand, explicitly uses the FBA solutions in its construction. Both methods provide a systematic framework to convert genome-scale metabolic models into a directed graph on which analysis tools from network theory can be applied.

To exemplify our approach, we built and analysed PFG and MFGs for the core metabolism of *E. coli*. Through the analysis of topological properties and community structure of these graphs, we highlighted the importance of weighted directionality in metabolic graph construction, and revealed the flux-mediated relationships between functional pathways under different environments. In particular, the MFGs capture specific metabolic adaptations such as the glycolytic-gluconeogenic switch, overflow metabolism, and the effects of anoxia. The proposed graph construction can be readily applied to large genome-scale metabolic networks [12, 19, 21, 22, 40].

To illustrate the scalability of our analyses to larger metabolic models, we studied a genome-scale model of a large metabolic model of human hepatocytes with around 3000 reactions in which we compared the wild type and a mutated state associated with the disease PH1 under more than 400 metabolic conditions [34]. Our network analysis of the MFGs revealed a consistent organisation of the reaction graph, which is highly preserved under the mutation. Our analysis also identified notable changes in the network centrality score and community structure of certain reactions, which is linked to key biological processes in PH1. Importantly, network measures computed from the MFGs reveal complementary information to that provided by the sole analysis of perturbed FBA fluxes.

Our flux graphs provide a systematic connection between network theory and constraint-based methods widely employed in metabolic modelling [21, 22, 25, 32], thus opening avenues towards environment-dependent, graph-based analyses of cell metabolism. An area of interest for future research is the use of MFGs to study how network measures of flux graphs can help characterise metabolic conditions that maximise the efficacy of drug treatments or disease-related distortions, e.g., cancer-related metabolic signatures [57–60]. In particular, MFGs can quantify metabolic robustness via graph statistics upon removal of reaction nodes [22].

The proposed graph construction framework can be extended in different directions. The core idea behind our framework is the distinction between production and consumption fluxes, and how to encode both in the links of a graph. This general principle can also be used to build other potentially useful graphs. For example, two other graphs that describe relationships between reactions are:

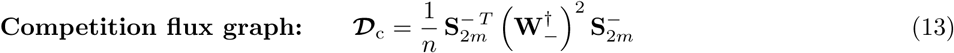

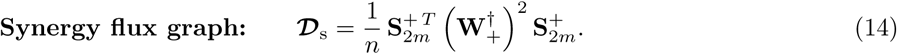

The competition and synergy graphs are undirected and their edge weights represent the probability that two reactions consume ( _c_) or produce ( _s_) metabolites chosen at random. The corresponding FBA versions of *competition and synergy flux graphs*, which follow directly from (12), (13) and (14) could help reveal further relationships between metabolic reactions in the cell. These graphs will be the subject of future studies.

Our approach could also be extended to include dynamic adaptations of metabolic activity: by using dynamic extensions of FBA [61, 62]; by incorporating static [63] or time-varying [64] enzyme concentrations; or by considering kinetic models (with kinetic constants when available) to generate probabilistic reactions fluxes in the sense of stochastic chemical kinetics [37, 65]. Of particular interest to metabolic modelling, we envision that MFGs could provide a novel route to evaluate the robustness of FBA solutions [25, 66] by exploiting the non-uniqueness of the MFG from each FBA solution in the space of graphs. Such results could enhance the interface between network science and metabolic analysis, allowing for the systematic exploration of the system-level organisation of metabolism in response to environmental constraints and disease states.

## Appendix A: Methods

### 1. Flux balance analysis

Flux Balance Analysis (FBA) [25, 32] is a widely-adopted approach to analyse metabolism and cellular growth. FBA calculates the reaction fluxes that optimise growth in specific biological contexts. The main hypothesis behind FBA is that cells adapt their metabolism to maximise growth in different biological conditions. The conditions are encoded as constraints on the fluxes of certain reactions; for example, constraints reactions that import nutrients and other necessary compounds from the exterior.

The mathematical formulation of the FBA is described in the following constrained optimisation problem:

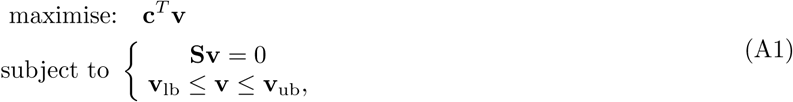

where **S** is the stoichiometry matrix of the model, **v** the vector of fluxes, **c** is an indicator vector (i.e., *c*(*i*) = 1 when *i* is the biomass reaction and zero everywhere else) so that **c**^*T*^ **v** is the flux of the biomass reaction. The constraint **Sv** = 0 enforces mass-conservation at stationarity, and **v**_lb_ and **v**_ub_ are the lower and upper bounds of each reaction’s flux. Through these vectors, one can encode a variety of different scenarios [33]. The biomass reaction represents the most widely-used flux that is optimised, although there are others can be used as well [31, 67].

In our simulations, we set the individual carbon intake rate to 18.5 mmol/gDW/h for every source available in each scenario. We allowed oxygen intake to reach the maximum needed in to consume all the carbon except in the anaerobic condition scenario, in which the upper bound for oxygen intake was 0 mmol/gDW/h. In the scenario with limited phosphate and ammonium intake, the levels of NH_4_ and phosphate intake were fixed at 4.5 mmol/gDW/h and 3.04 mmol/gDW/h respectively (a reduction of 50% compared to a glucose-fed aerobic scenario with no restrictions).

### 2. Markov Stability community detection framework

We extract the communities in each network using the Markov Stability community detection framework [50, 51]. This framework uses diffusion processes on the network to find groups of nodes (i.e., communities) that retain flows for longer than one would expect on a comparable random network; in addition, Markov Stability incorporates directed flows seamlessly into the analysis [49, 52].

The diffusion process we use is a continuous-time Markov process on the network. From the adjacency matrix **G** of the graph (in our case, the RAG, PFG or MFG), we construct a rate matrix for the process: 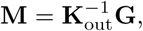, where out **K**_out_ is the diagonal matrix of out-strengths, *k*_out*,i*_ = _*j*_ *g*_*i,j*_. When a node has no outgoing edges then we simply let *k*_out*,i*_ = 1. In general, a directed network will not be strongly-connected and thus a Markov process on **M** will not have a unique steady state. To ensure the uniqueness of the steady state we must add a *teleportation* component to the dynamics by which a random walker visiting a node can follow an outgoing edge with probability λ or jump (teleport) uniformly to any other node in the network with probability 1 λ [43]. The rate matrix of a Markov process with teleportation is:

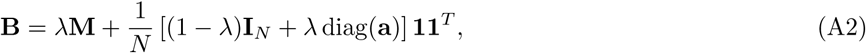

where the *N* × 1 vector **a** is an indicator for dangling nodes: if node *i* has no outgoing edges then *a_i_* = 1, and *a*_*i*_ = 0 otherwise. Here we use λ = 0.85. The Markov process is described by the ODE:

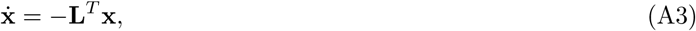

A hard partition of the graph into *C* communities can be encoded into the *N × C* matrix **H**, where *h_ic_* = 1 if node *i* belongs to community *c* and zero otherwise. The *C × C clustered autocovariance matrix* of (A3) is

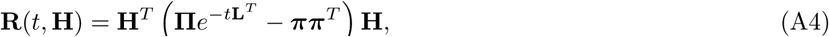

and the entry (*c, s*) of **R**(*t,* **H**) measures how likely it is that a random walker that started the process in community *c* finds itself in community *s* after time *t* when at stationarity. The diagonal elements of **R**(*t,* **H**) thus record how good the communities in **H** are at retaining flows. The *Markov stability of the partition* is then defined as

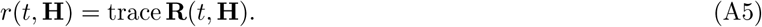

The optimised communities are obtained by maximising the cost function (A5) over the space of all partitions for every time *t* to obtain an optimised partition 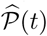. This optimisation is NP-hard; hence with no guarantees of optimality. Here we use the Louvain greedy optimisation heuristic [68], which is known to give high quality solutions 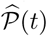 in an efficient manner. The value of the Markov time *t*, i.e. the duration of the Markov process, can be understood as a resolution parameter for the partition into communities [48, 50]. In the limit *t →* 0, Markov stability will assign each node to its own community; as *t* grows, we obtain larger communities because the random walkers have more time to explore the network [51]. We scan through a range of values of *t* to explore the multiscale community structure of the network. The code for Markov Stability can be found at github.com/michaelschaub/PartitionStability.

To identify the important partitions across time, we use two criteria of robustness [48]. Firstly, we optimise (A5) 100 times for each value of *t* and we assess the consistency of the solutions found. A relevant partition should be a robust outcome of the optimisation, i.e., the ensemble of optimised solutions should be similar as measured with the normalised variation of information [69]:

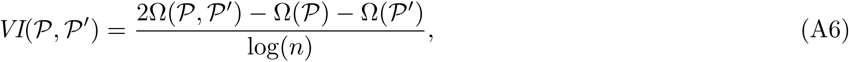

where Ω(*P*) = − *_C_ p*(*C*) log *p*(*C*) is a Shannon entropy and *p*(*C*) is the relative frequency of finding a node in community *C* in partition *P*. We then compute the average variation of information of the ensemble of solutions from the *f* = 100 Louvain optimisations *P_i_*(*t*) at each Markov time *t*:

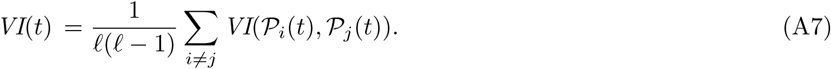

If all Louvain runs return similar partitions, then *VI*(*t*) is small, indicating robustness of the partition to the optimisation. Hence we select partitions with low values (or dips) of *VI*(*t*). Secondly, relevant partitions should also be optimal across Markov time, as indicated by a low values of the cross-time variation of information:

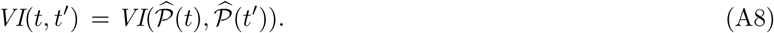

Therefore, we also search for partitions with extended low value plateaux of *VI*(*t, t^!^*) [48, 49, 55].

## ACKNOWLEDGMENTS

M.B.D. acknowledges support from the James S. McDonnell Foundation Postdoctoral Program in Complexity Science/Complex Systems Fellowship Award (#220020349-CS/PD Fellow), and the Oxford-Emirates Data Science Lab. G.B. acknowledges the support from the Spanish Ministry of Economy FPI Program (BES-2012-053772). D.O. acknowledges support from an Imperial College Research Fellowship and from the Human Frontier Science Program through a Young Investigator Grant (RGY0076-2015). J.P. acknowledges the support from the Spanish Ministry of Economy and EU FEDER through the SynBioFactory project (CICYT DPI2014-55276-C5-1). M.B. acknowledges funding from the EPSRC through grants EP/I017267/1 and EP/N014529/1.

*Data statement:* No new data was generated during the course of this research.

## Supplementary Information

## Appendix S1: Relation of the PFG with a directed version of the RAG

A directed version of the RAG (3) could in principle be obtained from the boolean production/consumption matrices 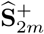 and 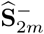 as follows. Projecting onto the space of reactions gives the 2*m* × 2*m* (asymmetric) adjacency matrix

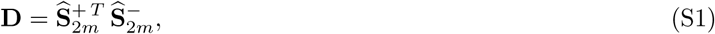

where the entries *D*_*ij*_ represent the total number of metabolites *produced* by reaction *R*_*i*_ that are *consumed* by reaction *R*_*j*_. A directed version of the Reaction Adjacency Graph on *m* nodes (directly comparable to the standard RAG) is then

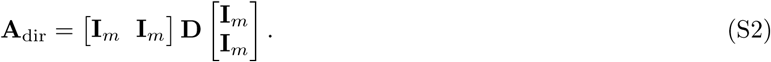

Clearly, when the metabolic model contains only reversible reactions, (i.e., the reversibility vector is all ones, 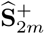, it follows that **A**_dir_ = **A**.

Although **A**_dir_ does not include spurious edges introduced by non-existent backward reactions, its structure is still obscured by the effect of uninformative connections created by pool metabolites.

## Appendix S2: Details of the toy metabolic network

As an illustration of the graph construction, the toy metabolic network in Fig. 1 was taken from Ref. [32]. The graph matrices for this model are as follows:

- Reaction Adjacency Graph, (Eq. 3):

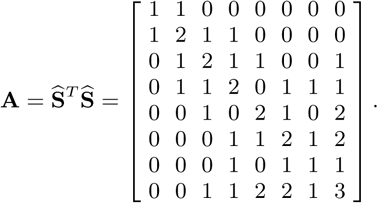
- Probabilistic Flux Graph, (Eq. 8):

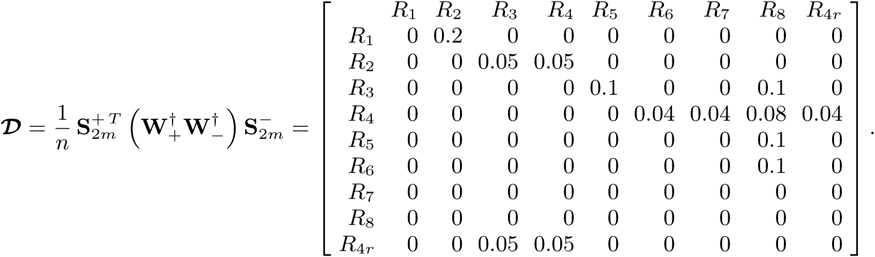
- Metabolic Flux Graph for FBA scenario 1, (Eq. 12):

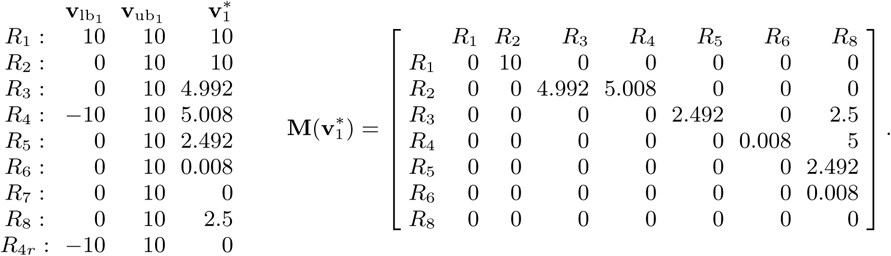
- Metabolic Flux Graph for FBA scenario 2, (Eq. 12):

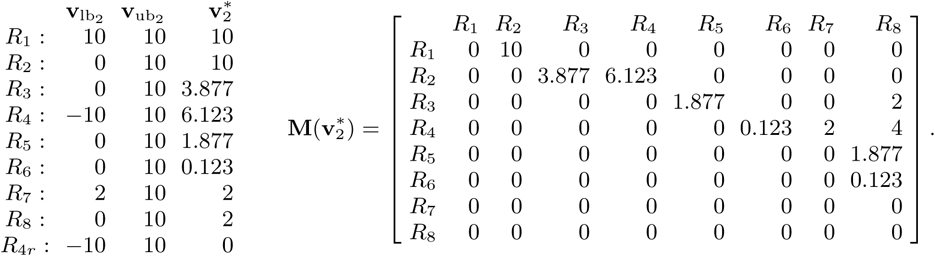

## Appendix S3: Reaction communities in context-free graphs of the core *E. coli* metabolic model

### 1. Reaction Adjacency Graph, A

A robust partition into seven communities in the RAG was found at Markov time *t* = 6.01 (Fig. S1A). The communities at this resolution (Fig. 3E) are:

- Community C1(**A**) contains all the reactions that consume or produce ATP and water (two pool metabolites). Production of ATP comes mostly from oxidative phosphorylation (ATPS4r) and substrate level phosphorylation reactions such as phosphofructokinase (PFK), phosphoglicerate kinase (PGK) and succinil-CoA synthase (SU-COAS). Reactions that consume ATP include glutamine synthetase (GLNS) and ATP maintenance equivalent reaction (ATPM). The reactions L-glutamine transport via ABC system (GLNabc), acetate transport in the form of phosphotransacetilase (PTAr), and acetate kinase (ACKr) are also part of this community. Additionally, C1(**A**) (green) contains also reactions that involve H_2_O. Under normal conditions water is assumed to be abundant in the cell, thus the biological link that groups these reactions together is tenuous.
- Community C2(**A**) includes the reactions NADH dehydrogenase (NADH16), cytochrome oxidase (CYTBD), and transport and exchange reactions. These two reactions involve pool metabolites (such as H^+^) which create a large number of connection. Other members include fumarate reductase (FR7) and succinate dehydrogenase (SUCDi) which couple the TCA cycle with the electron transport chain (through ubiquinone-8 reduction and ubiquinol-8 oxidation). Reactions that include export and transport of most secondary carbon sources (such as pyruvate, ethanol, lactate, acetate, malate, fumarate, succinate or glutamate) are included in the community as well. These reactions are included in the community because of their influence in the proton balance of the cell. Most of these reactions do not occur under normal circumstances. This community highlights the fact that in the absence of biological context, many reactions that do not normally interact can be grouped together.
- Community C3(**A**) contains reactions that produce or consume nicotinamide adenine dinucleotide (NAD^+^), nicotinamide adenine dinucleotide phosphate (NADP^+^), or their reduced variants NADH and NADPH. The main two reactions of the community are NAD(P) transhydrogenase (THD2) and NAD^+^ transhydrogenase (NADTRHD). There are also reactions related to the production of NADH or NADPH in the TCA cycle such as isocitrate dehydrogenase (ICDHyr), 2-oxoglutarate dehydrogenase (AKGDH) and malate dehydrogenase (MDH). The community also includes reactions that are not frequently active such as malic enzime NAD (ME1) and malic enzime NADH (ME2) or acetate dehydrogenase (ACALD) and ethanol dehydrogenase (ALCD2x).
- Community C4(**A**) contains the main carbon intake of the cell (glucose), the initial steps of glycolysis, and most of the pentose phosphate shunt. These reactions are found in this community because the metabolites involved in these reactions (e.g., alpha-D-ribose-5-phosphate (r5p) or D-erythrose-4-phosphate (e4p)) are only found in these reactions. This community includes the biomass reaction due to the number of connections created by growth precursors.
- Communities C5(**A**), C6(**A**) and C7(**A**) are small communities that contain oxygen intake, ammonium intake and acetaldehyde secretion reactions, respectively.

**FIG. S1.**
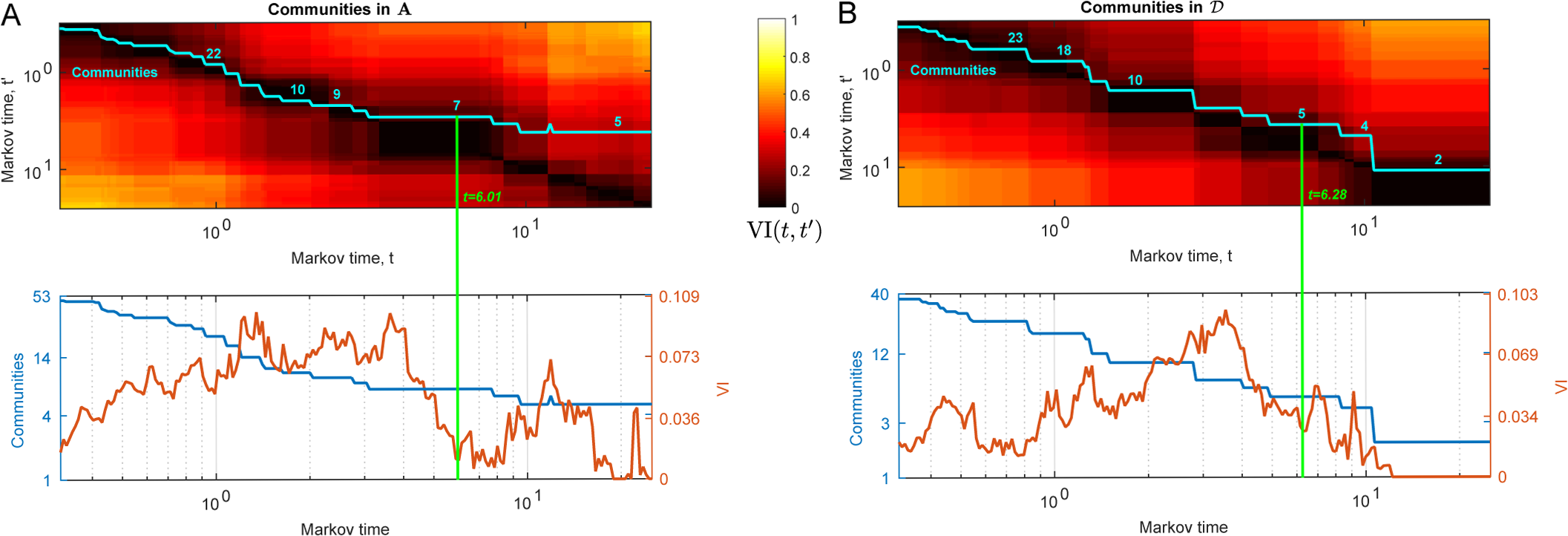
Community structure in the template networks A and **𝒟**. (A) Communities in **A**. Top plot: Variation of Information (VI) of the best partition found at Markov time *t* with every other partition at time *t^!^*. Bottom plot: Number of communities and VI of the ensemble of solutions found at each Markov time. A robust partition into seven communities is found at *t* = 6.01. (B) Communities and VI in **𝒟**. A robust partition into five communities is found at *t* = 6.28.

### 2. Probabilistic Flux Graph, **𝒟**

A robust partition into five communities in the PFG was found at Markov time *t* = 6.28 (Fig. S1B). The communities at this resolution (Fig. 3C) are:

- Community C1(**𝒟**) includes the first half of the glycolysis and the complete pentose phosphate pathway. The metabolites that create the connections among these reactions such as D-fructose, D-glucose, or D-ribulose.
- Community C2(**𝒟**) contains the main reaction that produces ATP through substrate level (PGK, PYK, ACKr) and oxidative phosphorylation (ATPS4r). The flow of metabolites among the reactions in this community includes some pool metabolites such as ATP, ADP, H_2_0, and phosphate. However, there are connections created by metabolites that only appear in a handful of reactions such as adenosine monophosphate (AMP) whose sole producer is phosphoenolpyruvate synthase (PPS) and its sole consumer is ATPS4r. This community also contains the biomass reaction.
- Community C3(**𝒟**) includes the core of the citric acid (TCA) cycle such as citrate synthase (CS), aconitase A/B (ACONTa/b), and anaplerotic reactions such as malate synthase (MALS), malic enzyme NAD (ME1), and malic enzyme NADP (ME2). This community also includes the intake of cofactors such as CO_2_.
- Community C4(**𝒟**) contains reactions that are secondary sources of carbon such as malate and succinate, as well as oxidative phosphorilation reactions.
- Community C5(**𝒟**) contains some reactions part of the pyruvate metabolism subsystem such as D-lactate dehydrogenase (LDH-D), pyruvate formate lyase (PFL) or acetaldehyde dehydrogenase (ACALD). In addition, it also includes the tranport reaction for the most common secondary carbon metabolites such as lactate, formate, acetaldehyde and ethanol.

## Appendix S4 Reaction communities in Metabolic Flux Graphs of *E. coli* metabolism under different biological scenarios

### 1. M_glc_: aerobic growth under glucose

This graph has 48 reactions with nonzero flux and 227 edges. At Markov time *t* = 7.66 (Fig. S2A) this graph has a partition into three communities (Fig. 4A):

- Community C1(**M**_glc_) comprises the intake of glucose and most of the glycolysis and pentose phosphate pathway. The function of the reactions in this community consists of carbon intake and processing glucose into phosphoenolpyruvate (PEP). This community produces essential biocomponents for the cell such as alpha-D-Ribose 5-phosphate (rp5), D-Erythrose 4-phosphate (e4p), D-fructose-6-phosphate (f6p), glyceraldehyde-3-phosphate (g3p) or 3-phospho-D-glycerate (3pg). Other reactions produce energy ATP and have reductive capabilities for catabolism.
- Community C2(**M**_glc_) contains the electron transport chain which produces the majority of the energy of the cell. In the core *E coli* metabolic model the chain is represented by the reactions NADH dehydrogenase (NADH16), cytochrome oxidase BD (CYTBD) and ATP synthase (ATPS4r). This community also contains associated reactions to the electron transport such as phosphate intake (EXpi(e), PIt2), oxygen intake (EXo2(e), O2t) and proton balance (EXh(e)). This community also includes the two reactions that represent energy maintenance costs (ATPM), and growth (biomass); this is consistent with the biological scenario because ATP is the main substrate for both ATPM, and the biomass reaction.
- Community C3(**M**_glc_) contains the TCA cycle at its core. The reactions in this community convert PEP into ATP, NADH and NADPH. In contrast with C1(**M**_glc_), there is no precursor formation here. Beyond the TCA cycle, pyruvate kinase (PYK), phosphoenolpyruvate carboxylase (PPC) and pyruvate dehydrogenase (PDH) appear in this community. These reactions highlight the two main carbon intake routes in the cycle: oxalacetate from PEP through phosphoenol pyruvate carboxylase (PPC), and citrate from acetyl coenzyme A (acetyl-CoA) via citrate synthase (CS). Furthermore, both routes begin with PEP, so it is natural for them to belong to the same community along with the rest of the TCA cycle. Likewise, the production of L-glutamate from 2-oxoglutarate (AKG) by glutamate dehydrogenase (GLUDy) is strongly coupled to the TCA cycle.

### 2. M_etoh_: aerobic growth under ethanol

This graph contains 49 reactions and 226 edges. At Markov time *t* = 6.28 (Fig. S2B) this graph has a partition into three communities (Fig. 4B):

- Community C1(**M**_etoh_) in this graph is similar to its counterpart in **M**_glc_, but with important differences. For example, the reactions in charge of the glucose intake (EXglc(e) and GLCpts) are no longer part of the network (i.e., they have zero flux), and reactions such as malic enzyme NAPD (ME2) and phosphoenolpyruvate caboxyk-inase (PPCK), which now appear in the network, belong to this community. This change in the network reflect the cell’s response to a new biological situation. The carbon intake through ethanol has changed the direction of glycolysis into gluconeogenesis [1] (the reactions in C1(**M**_glc_) in Fig. 4A are now operating in the reverse direction in Fig. 4B). The main role of the reactions in this community is the production of bioprecursors such as PEP, pyruvate, 3-phospho-D-glycerate (3PG) glyceraldehyde-3-phosphate (G3P), D-fructose-6-phosphate (F6P), and D-glucose-6-phosphate, all of which are substrates for growth. Reactions ME2 and PPCK also belong to this community due to their production of PYR and PEP. Reactions that were in a different community in **M**_glc_, such as GLUDy and ICDHyr which produce precursors L-glutamate and NADPH respectively, are now part of C1(**M**_etoh_). This community also includes the reactions that produce inorganic substrates of growth such as NH_4_, CO_2_ and H_2_O.
- Community C2(**M**_etoh_) contains the electron transport chain and the bulk of ATP production, which is similar to C2(**M**_glc_). However, there are subtle differences that reflect changes in this new scenario. Ethanol intake and transport reactions (EXetoh(e) and ETOHt2r) appear in this community due to their influence in the proton balance of the cell. In addition, C2(**M**_etoh_) contains NADP transhydrogenase (THD2) which is in charge of NADH/NADPH balance. This reaction is present here due to the NAD consumption involved in the reactions ACALD and ethanol dehydrogenase (ALCD2x), which belong to this community as well.
- Community C3(**M**_etoh_) contains most of the TCA cycle. The main difference between this community and C1(**M**_glc_) is that here acetyl-CoA is extracted from acetaldehyde (which comes from ethanol) by the reaction acetaldehyde dehydrogenase reaction (ACALD), instead of the classical pyruvate from glycolysis. The glycoxy-late cycle reactions isocitrate lyase (ICL) and malate synthase (MALS) which now appear in the network, also belong to this community. These reactions are tightly linked to the TCA cycle and appear when the carbon intake is acetate or ethanol to prevent the loss of carbon as CO_2_.

### 3. M_anaero_: anaerobic growth

This graph contains 47 reactions and 212 edges. At Markov time *t* = 6.01 (Fig. S2C) this graph has a partition into four communities (Fig. 4C):

- Community C1(**M**_anaero_) contains the reactions responsible D-glucose intake (EXglc) and most of the glycolysis. The reaction that represents the cellular maintenance energy cost, ATP maintenance requirement (ATPM), is included in this community because of the increased strength of its connection to the substrate-level phosphori-lation reaction phosphoglycerate kinase (PGK). Also note that reactions in the pentose phosphate pathway do not belong to the same community as the glycolysis reactions (unlike in **M**_glc_ and **M**_etoh_).
- Community C2(**M**_anaero_) contains the conversion of PEP into formate through the sequence of reactions PYK, PFL, FORti and EXfor(e). More than half of the carbon secreted by the cell becomes formate.
- Community C3(**M**_anaero_) includes the biomass reaction and the reactions in charge of supplying it with substrates. These reactions include the pentose phosphate pathway (now detached from C1(**M**_glc_)), which produce essential growth precursors such as alpha-D-ribose-5-phosphate (r5p) or D-erythrose-4-phosphate (e4p). The TCA cycle is present as well because its production of two growth precursors: 2-oxalacetate and NADPH. Finally, the reactions in charge of acetate production (ACKr, ACt2r and EXac(e)) are also members of this community through the ability of ACKr to produce ATP. Glutamate metabolism reaction GLUDy is also included in this community. It is worth mentioning that the reverse of ATP synthase (ATPS4r) is present in this community because here, unlike in **M**_glc_, ATPS4r consumes ATP instead of producing it. When this flux is reversed, then ATPS4r is in part responsible for pH homeostasis.
- Community C4(**M**_anaero_) includes the main reactions involved in NADH production and consumption, which occurs via glyceraldehyde-3-phosphate dehydrogenase (GAPD). NADH consumption occurs in two consecutive steps in ethanol production: in ACALD and ALCD2x. The phosphate intake and transport reactions EXpi(e) and PIt2r belong to this community because most of the phosphate consumption takes place at GAPD. Interestingly, the core reaction around which the community forms (GAPD) is not present in the community. It is included in earlier Markov times but when communities start to get larger the role of GAPD becomes more relevant as a part of the glycolysis than its role as a NADH hub. This is a good example of how the graph structure and the clustering method are able to capture two different roles in the same metabolite.

### 4. M_lim_: aerobic growth under limiting conditions

This graph has 52 nodes and 228 edges. At Markov time *t* = 13 this graph (Fig. S2D) has a partition into three communities (Fig. 4D):

- Community C1(**M**_lim_) contains the glycolysis pathway (detached from the pentose phosphate pathway). This community is involved in precursor formation, ATP production, substrate-level phosphorylation and processing of D-glucose into PEP. Community C2(**M**_lim_) contains the bioenergetic machinery of the cell; the main difference to the previous scenarios is that the electron transport chain has a smaller role in ATP production (ATPS4r), and substrate-level phosphorylation (PGK, PYK, SUCOAS, ACKr) becomes more important. In **M**_lim_ the electron transport chain is responsible for the 21.8% of the total ATP produced in the cell while in **M**_glc_ it produces 66.5%. The reactions in charge of intake and transport of inorganic ions such as phosphate (EXpi(e) and PIt2r), O_2_ (EXO_2_(e) and O_2_t)and H_2_O (EXH_2_O and H_2_Ot) belong to this community as well. This community includes the reactions in the pentose phosphate pathway that produce precursors for growth: transketolase (TKT2) produces e4p, and ribose-5-phosphate isomerase (RPI) produces r5p.
- Community C3(**M**_lim_) is the community that differs the most from those in the other aerobic growth networks (**M**_glc_ and **M**_etoh_). This community gathers reactions that under normal circumstances would not be so strongly related but that the limited availability of ammonium and phosphate have forced together; its members include reactions from the TCA cycle, the pentose phosphate pathway, nitrogen metabolism and by-product secretion. The core feature of the community is carbon secretion as formate and acetate. Reactions PPC, malate dehydrogenase (MDH) reverse and ME2 channel most of the carbon to the secretion routes in the form of formate and acetate. The production of L-glutamine seems to be attached to this subsystem through the production of NADPH in ME2 and its consumption in the glutamate dehydrogenase NAPD (GLUDy).

**FIG. S2.**
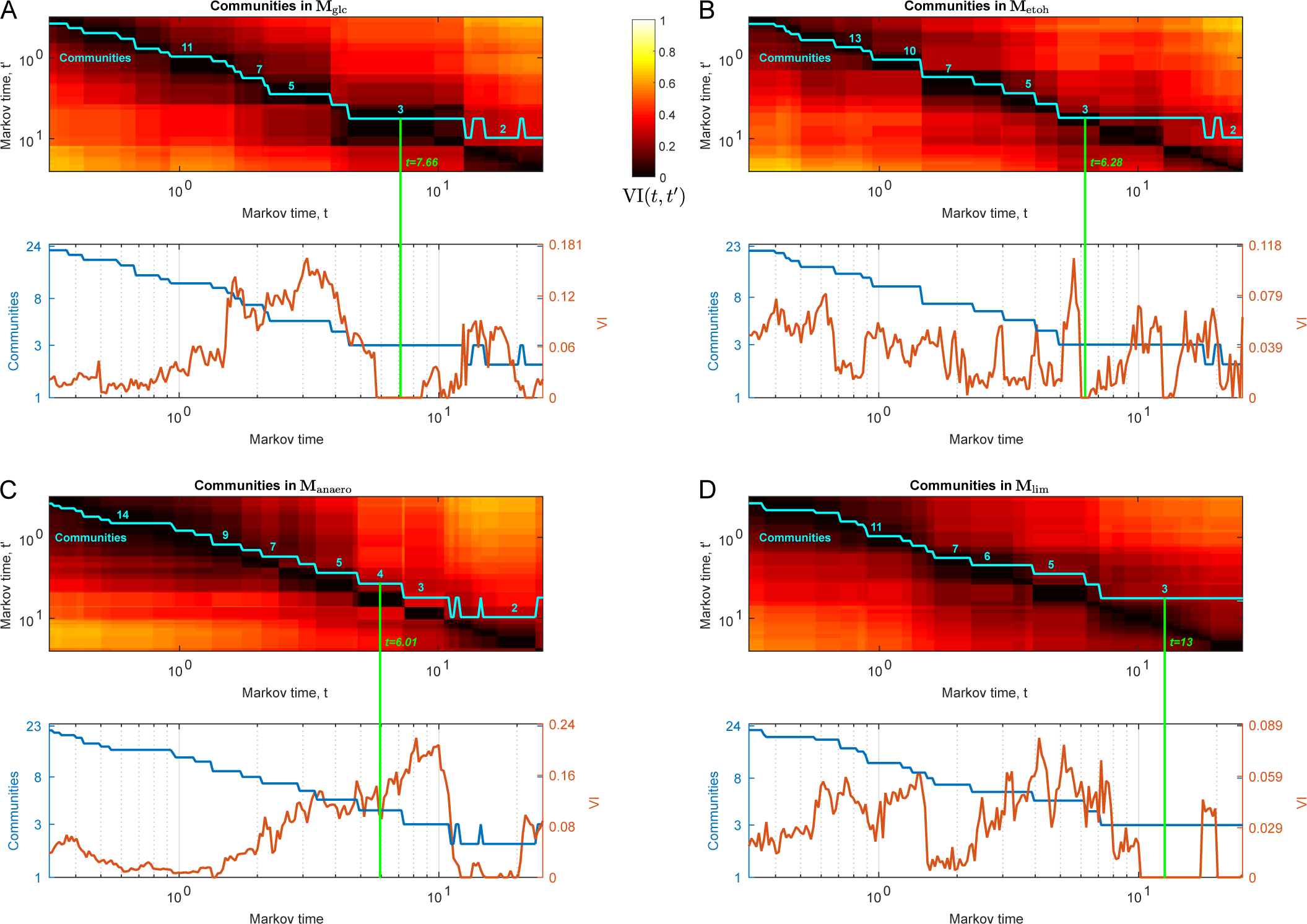
Community structure in the MFGs. Number of communities and VI of the MFGs in four biological scenarios. (A) The graph **M**_glc_ has a robust partition into three communities at *t* = 7.66. (B) **M**_etoh_ has a partition into three communities at *t* = 6.28. (C) **M**_anaero_ has four communities at *t* = 6.01. (D) **M**_lim_ has three communities at *t* = 13.0.

